# Artificial Intelligence Guided Discovery of Gastric Cancer Continuum

**DOI:** 10.1101/2022.10.05.510975

**Authors:** Daniella Vo, Pradipta Ghosh, Debashis Sahoo

## Abstract

**Background:** Detailed understanding of pre, early and late neoplastic states in gastric cancer helps develop better models of risk of progression to Gastric Cancers (GCs) and medical treatment to intercept such progression.

**Methods:** We built a Boolean Implication network of gastric cancer and deployed machine learning algorithms to develop predictive models of known pre-neoplastic states, e.g., atrophic gastritis, intestinal metaplasia (IM) and low-to high-grade intestinal neoplasia (L/HGIN), and GC. Our approach exploits the presence of asymmetric Boolean Implication relationships that are likely to be invariant across almost all gastric cancer datasets. Invariant asymmetric Boolean Implication relationships can decipher fundamental time series underlying the biological data. Pursuing this method, we developed a healthy mucosa →GC continuum model based on this approach.

**Results:** Our model performed better against publicly available models for distinguishing healthy versus GC samples. Although not trained on IM and L/HGIN datasets, the model could identify the risk of progression to GC via the metaplasia →dysplasia →neoplasia cascade in patient samples. The model could rank all publicly available mouse models for their ability to best recapitulate the gene expression patterns during human GC initiation and progression.

**Conclusions:** A Boolean implication network enabled the identification of hitherto undefined continuum states during GC initiation. The developed model could now serve as a starting point for rationalizing candidate therapeutic targets to intercept GC progression.

**MINI-ABSTRACT:** We developed predictive models of early and late neoplastic states in gastric cancer and identified gene clusters that are up/down-regulated at various points along the gastric cancer disease continuum.

## INTRODUCTION

Gastric cancer (GC) often presents as an advanced disease with patients either having inoperable conditions or surgery as the only potentially curative treatment [1]. There is evidence that 75% of all GCs are initiated by *Helicobacter pylori*, a known carcinogenic pathogen [2, 3]. Risk factors also include age, sex, smoking and family history [4]. This oncogenesis leads to Correa’s cascade, a stepwise progression from normal, chronic active gastritis, atrophic gastritis, intestinal metaplasia, dysplasia then adenocarcinomas [3]. Intestinal metaplasia also has two subtypes, incomplete and complete intestinal metaplasia (IIM and CIM, respectively), with IIM having a higher probability of developing GC compared to CIM [5].

Research into GCs has used impactful approaches to investigate the genome [6], therapeutics [7] and survival [8], but these methods have not translated into actionable biomarkers of prognostication, targets, novel therapeutics, or changes in screening strategies. These genomic insights also have not provided insight into which genes are important in the progression of GC for preneoplastic detection and treatment.

Here we present a network-based approach for biomarker and target discovery that uses artificial intelligence (AI) to select genes and then perform rigorous validation in multiple independent GC datasets. Previously, we have successfully exploited this approach to identify biomarkers in IBD [9], COVID-19 [10] and macrophages [11]. We demonstrate how Boolean implications allow us to develop models that provide insight into the gastric cancer disease continuum.

## METHODS

Detailed methods for computational modeling and AI-guided target identification are presented in Online Resource 1 and mentioned in brief here.

### Construction of a Network of Boolean Implications

Modeling continuum states within the metaplasia → dysplasia → neoplasia cascade was performed using Boolean Network Explorer (BoNE) [9]. We created an asymmetric gene expression network, for the progression from normal to gastric cancer (GC), using a computational method based on Boolean logic [12]. To build the GC network, we analyzed a publicly available gastric cancer transcriptomic dataset, GSE66229[13] (n = 400; 300 GC tumor and 100 patient-matched normal tissue). A Boolean Network Explorer (BoNE; see **Online Resource 1** for more details) computational tool was introduced, which uses asymmetric properties of Boolean implication relationships (BIRs as in MiDReG algorithm [12]) to model natural progressive time-series changes in major cellular compartments that initiate, propagate, and perpetuate cellular state change and are likely to be important for GC progression. BoNE provides an integrated platform for the construction, visualization and querying of a network of progressive changes much like a disease map (in this case, GC map) in three steps: First, the expression levels of all genes in these datasets were converted to binary values (high or low) using the StepMiner algorithm [14]. Second, gene expression relationships between pairs of genes were classified into one-of-six possible BIRs and expressed as Boolean implication statements; two symmetric Boolean implications “equivalent” and “opposite” are discovered when two diagonally opposite sparse quadrants are identified and four asymmetric relationships, each corresponding to one sparse quadrant. While conventional symmetric analysis of transcriptomic datasets can recognize the latter 2 relationships, such an approach ignores the former. BooleanNet statistics are used to assess the significance of the Boolean implication relationships [12]. Prior work [9] has revealed how the Boolean approach offers a distinct advantage from currently used conventional computational methods that rely exclusively on symmetric linear relationships from gene expression data, e.g., differential, correlation-network, coexpression-network, mutual information-network, and the Bayesian approach. The other advantage of using BIRs is that they are robust to the noise of sample heterogeneity (i.e., healthy, diseased, genotypic, phenotypic, ethnic, interventions, disease severity) and every sample follows the same mathematical equation, and hence is likely to be reproducible in independent validation datasets. Third, genes with similar expression architectures, determined by sharing at least half of the equivalences among gene pairs, were grouped into clusters, and organized into a network by determining the overwhelming Boolean relationships observed between any two clusters. In the resultant Boolean implication network, clusters of genes are the nodes, and the BIR between the clusters are the directed edges; BoNE enables their discovery in an unsupervised way while remaining agnostic to the sample type. All gene expression datasets were visualized using Hierarchical Exploration of Gene Expression Microarrays Online (HEGEMON) framework [9].

### Ordering samples based on composite score of Boolean path

A Boolean path contains one or more clusters. A composite score is computed for each cluster and combined later. To compute the final score, first the genes present in each cluster were normalized and averaged. Gene expression values were normalized according to a modified Z-score approach centered around StepMiner threshold (formula = (expr - SThr)/3/stddev). A weighted linear combination of the averages from the clusters of a Boolean path was used to create a score for each sample. The weights along the path either monotonically increased or decreased to make the sample order consistent with the logical order based on BIR. The samples were ordered based on the final weighted and linearly combined score. A cluster highly expressed in a disease setting received a positive weight (ex: 1, 2, 3, etc.) and healthy setting received a negative weight (ex: -1, -2, -3, etc.).

### Multivariate Analysis for Model Selection

Two microarray datasets (GSE37023 (only samples on GPL96 Affymetrix Human Genome U133A Array used for analysis), n = 65, non-malignant = 36, GC tumor = 29; GSE122401, n = 160, patient-matched normal = 80, GC tumor = 80) are used to train a network model to distinguish normal vs GC samples. Using Ordinary Least Squares (OLS) regression in Python statsmodels (version 0.12.2), we performed multivariate analysis to determine which models performed best in the two training datasets.

### Statistical Analysis

Statistical significance between experimental groups was determined using Python scipy.stats.ttest_ind package (version 0.19.0) with Welch’s Two Sample t-test (two-tailed, unpaired, unequal variance (equal_var=False), and unequal sample size). For all tests, a p-value of 0.05 was used as the cutoff to determine significance. Violin and bar plots are created using Python seaborn package version 0.10.1.

## RESULTS

### Machine learning identified two possible Boolean paths in the GC disease map

Using a publicly available GC dataset (GSE66229) with tumor (T) and adjacent normal (AN) samples, we built a Boolean implication network (See *Methods* and **Online Resource 1**; **Fig. 1a**). Each cluster was evaluated to determine whether they fall on the healthy versus GC side of the disease map based on whether the average gene expression value of a cluster in healthy samples is up or down, yielding a GC map (**Fig. 1b**). We then used machine learning to identify Boolean paths (clusters connected by Boolean implication relationships) in the GC map that can distinguish tumor from AN samples in the training datasets (Figure 1C *top graphic*). Clusters #11-2-4-14 (C#11-2-4-14) performed the best with an ROC-AUC of 0.96 in training dataset #1 (GSE37023 AN versus T), while clusters #7-13-14 (C#7-13-14) performed best in training dataset #2 (GSE122401 AN vs T) with an ROC-AUC of 0.98 (**Fig. 1c**). Specific violin plots for both datasets and Boolean paths are presented in **Fig. 1d**. We performed Reactome pathway analysis on clusters in both paths to identify the top five biological processes associated with the clusters (**Fig. 1e**). Cluster 11 involves the downregulation of genes related to muscle contraction in GC. Cluster 2 represents genes relevant to cell cycle as many other studies pointed out their relevance in the context of GC [15, 16]. Cluster 4 had genes from the immune system including neutrophil degranulation as linked in other papers [17, 18]. Clusters 7 and 13 had genes involved in the downregulation of ion channel transport in GC [19, 20]. Cluster 14 represents genes increased in extracellular matrix processes [21, 22]. Since both Boolean paths C#11-2-4-14 and C#7-13-14 can distinguish AN versus GC samples, we identified a gene signature called GC-BoNE uses the path that best characterized the different samples (highest ROC-AUC score out of both paths) for classification of samples.

**Fig. 1.**
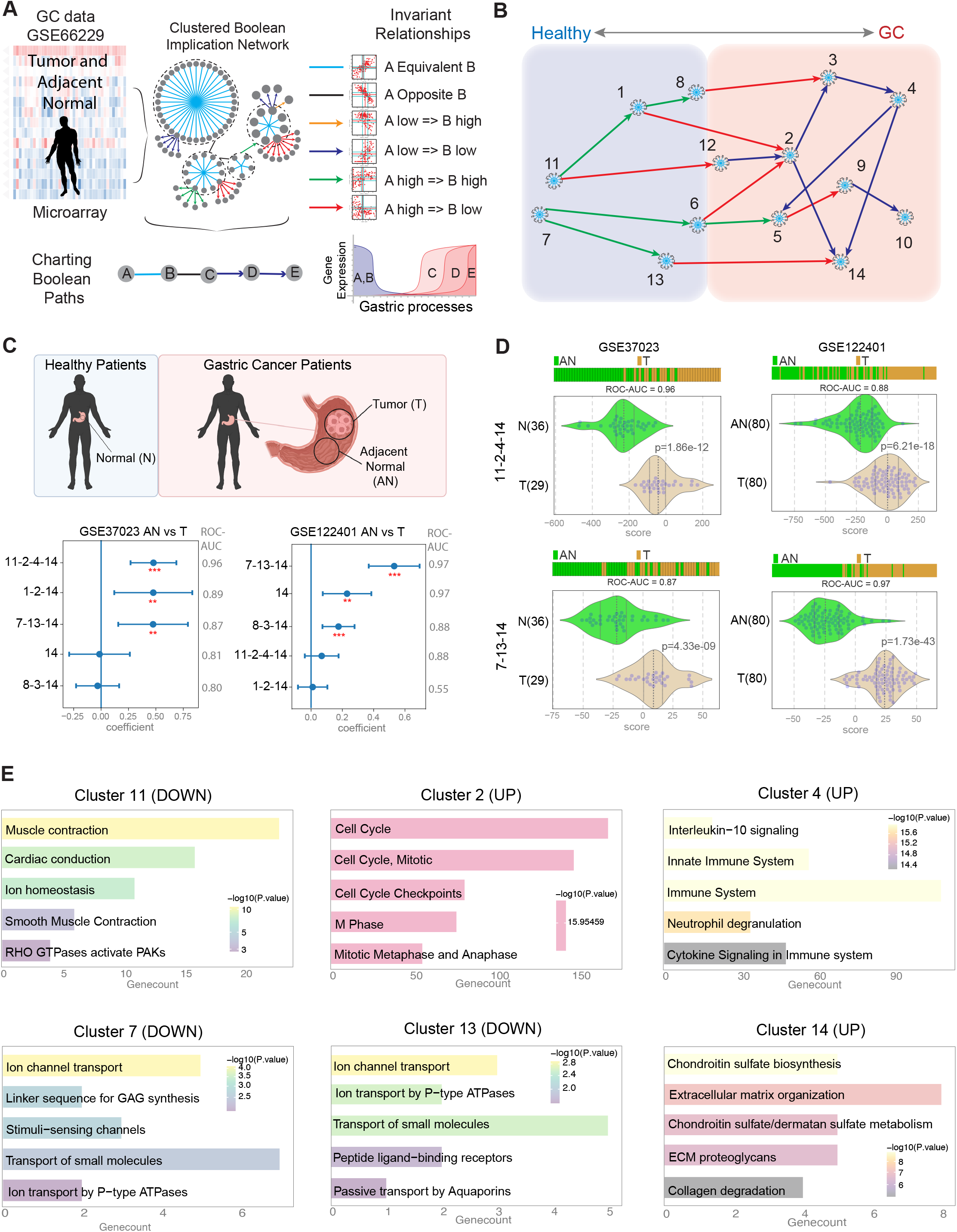
Generation and validation of Boolean implication network-derived gastric cancer (GC) signature. **a**. Schematic summarizing the workflow to build a Boolean map using a gastric cancer microarray dataset containing tumor and adjacent normal samples (GSE66229) **b**. Disease map representing the continuum from normal stomach to gastric cancer **c**. Selection of Boolean path using machine learning on two training datasets (GSE37023 and GSE122401). Multivariate regression was used to determine which path best separated the tumor from the adjacent normal samples. Coefficient of each path score (at the center) with 95% confidence intervals (as error bars) and the p values were illustrated in the bar plot. The p value for each term tests the null hypothesis that the coefficient is equal to zero (no effect) **d**. Violin plots showing the top Boolean paths in each of the training datasets **e**. Reactome pathway analysis of the gene clusters in the GC-BoNE signature

We tested how well the clusters identified by our Boolean approach would compare to previously established gene signatures (**Fig. 2a**). C#11-2-4-14 and C#7-13-14 individually (**Fig. 2b**) could classify the tumor and normal/adjacent normal samples in the 21 validation datasets (see **Online Resource 2** for a list of GSE IDs; ROC-AUC ranges from 0.57 - 1.00 in C#11-2-4-14, and 0.66 - 1.00 in C#7-13-14). We then compared GC-BoNE to other gene signatures (see **Online Resource 3** for list of genes in signatures; **Fig. 2c**) and found that our signature outperformed the others (average ROC-AUC for GC-BoNE is 0.933, and other signatures range from 0.690 - 0.921). There were minimal overlaps between clusters 11-2-4 (**Fig. 2d**), 7-13 (**Fig. 2e**) and the top three signatures (DEA (Li 2015), DEA+PPIN and Japanese GC). Cluster 14 and the Japanese GC signature had 8 overlapping genes (**Fig. 2f**). These findings suggest GC-BoNE provides a new list of potential biomarkers for GC that differ from previous signatures.

**Fig. 2.**
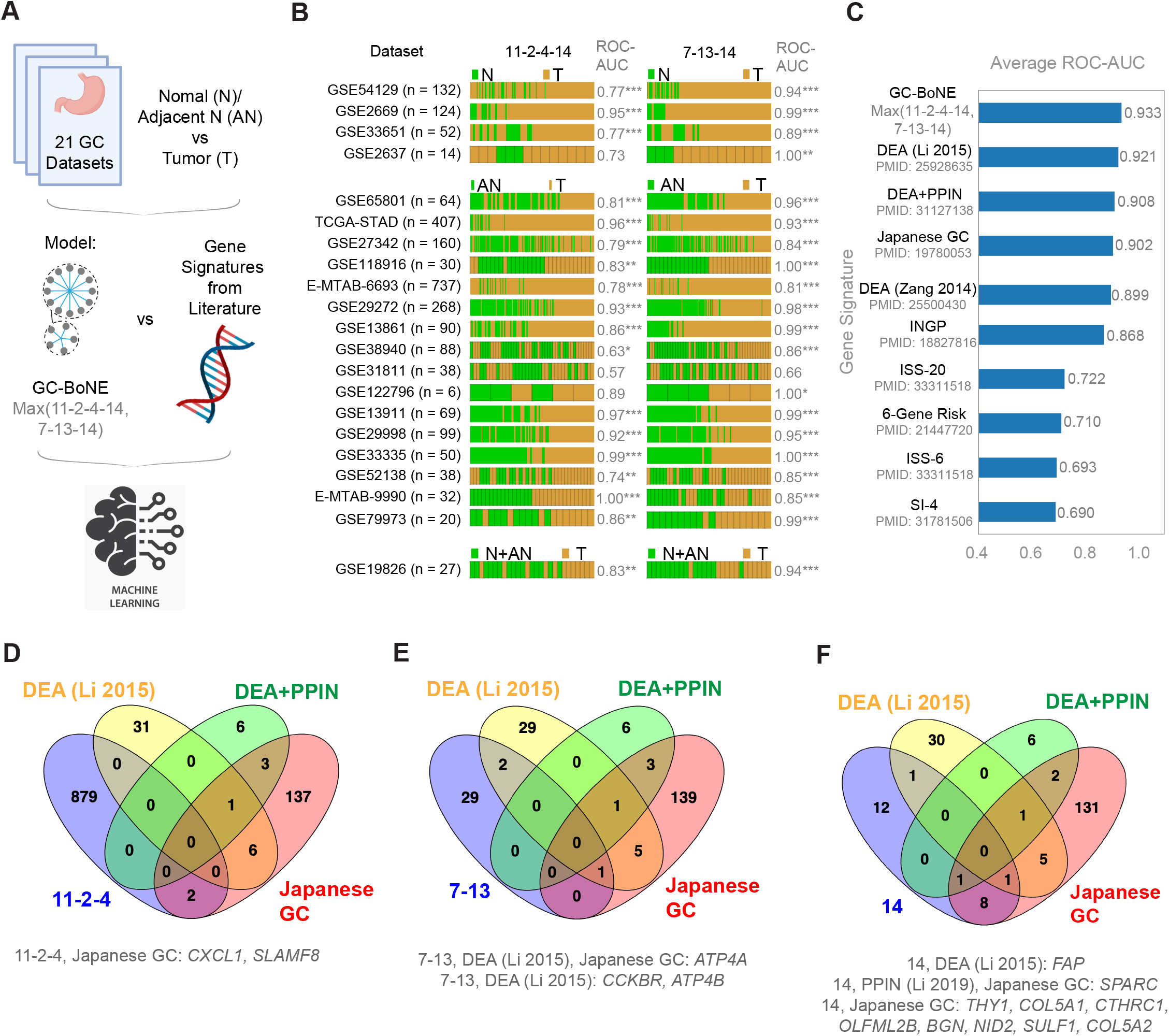
Comparison of classification accuracy using GC-BoNE signature versus gene signatures from previous literature for normal versus GC samples. **a**. Schematic summarizing the workflow to compare GC-BoNE to other gene signatures **b**. Bar plots of GC datasets comparing normal (N)/AN vs T showing the ROC-AUC values for the Boolean paths in the GC-BoNE signature (11-2-4-14 and 7-13-14). Asterisks (*) after the ROC-AUC values represent the following: *p<=0.05, **p<=0.01, ***p<=0.001, no asterisk: p-value>0.05 **c**. Comparison of average ROC-AUC values for the datasets in B. using GC-BoNE and other gene signatures [6, 48-54] (DEA: Differential Expression Analysis, PPIN: Protein-Protein Interaction Network, INGP: Ingenuity Pathway analysis, ISS: Immune Scoring System, SI: Stromal-Immune score; See Online Resource 3 for the complete list of genes in these signatures) **d-f**. Venn diagrams showing the overlaps in genes in the top four gene signatures (GC-BoNE, DEA (Li 2015), DEA+PPIN and Japanese GC)

### GC-BoNE identifies progressively increasing risk of GC along the metaplasia-dysplasia continuum

We next asked if the GC-BoNE signature is induced during the progression from normal to GC through the normal →inflammation (gastritis) →metaplasia →dysplasia →neoplasia cascade. In one dataset (E-MTAB-8889), we looked at the normal →inflammation (gastritis) →metaplasia cascade by comparing pairwise each sequential step, i.e., non-atrophic gastritis (NAG) vs chronic active gastritis (CG), CG vs chronic atrophic gastritis (CAG) and CAG vs intestinal metaplasia (IM) (**Fig. 3a**). We also looked at the first step in the cascade vs the other steps, i.e., NAG vs CAG and NAG vs IM (**Fig. 3a**). In another dataset (GSE55696), we studied the dysplasia →neoplasia cascade, which is typically scored by histopathological examination, as per the Vienna classification [23]; the latter comprises a continuum extending from low to high grade dysplasia to intramucosal carcinoma. Here, we looked at chronic gastritis (CG) vs low-grade intestinal neoplasia (LGIN), LGIN vs high-grade intestinal neoplasia (HGIN), HGIN vs early gastric cancer (EGC), CG vs HGIN and CG vs EGC (**Fig. 3b**). We compared GC-BoNE to the other signatures (**Fig. 3c**) and found that our signature again outperformed the others when looking at progression (see **Online Resource 2** for a list of GSE IDs; average ROC-AUC for GC-BoNE is 0.828, and other signatures range from 0.633 - 0.806). These findings suggest the genes identified in GC-BoNE may provide further insight into what initiates GC progression.

**Fig. 3.**
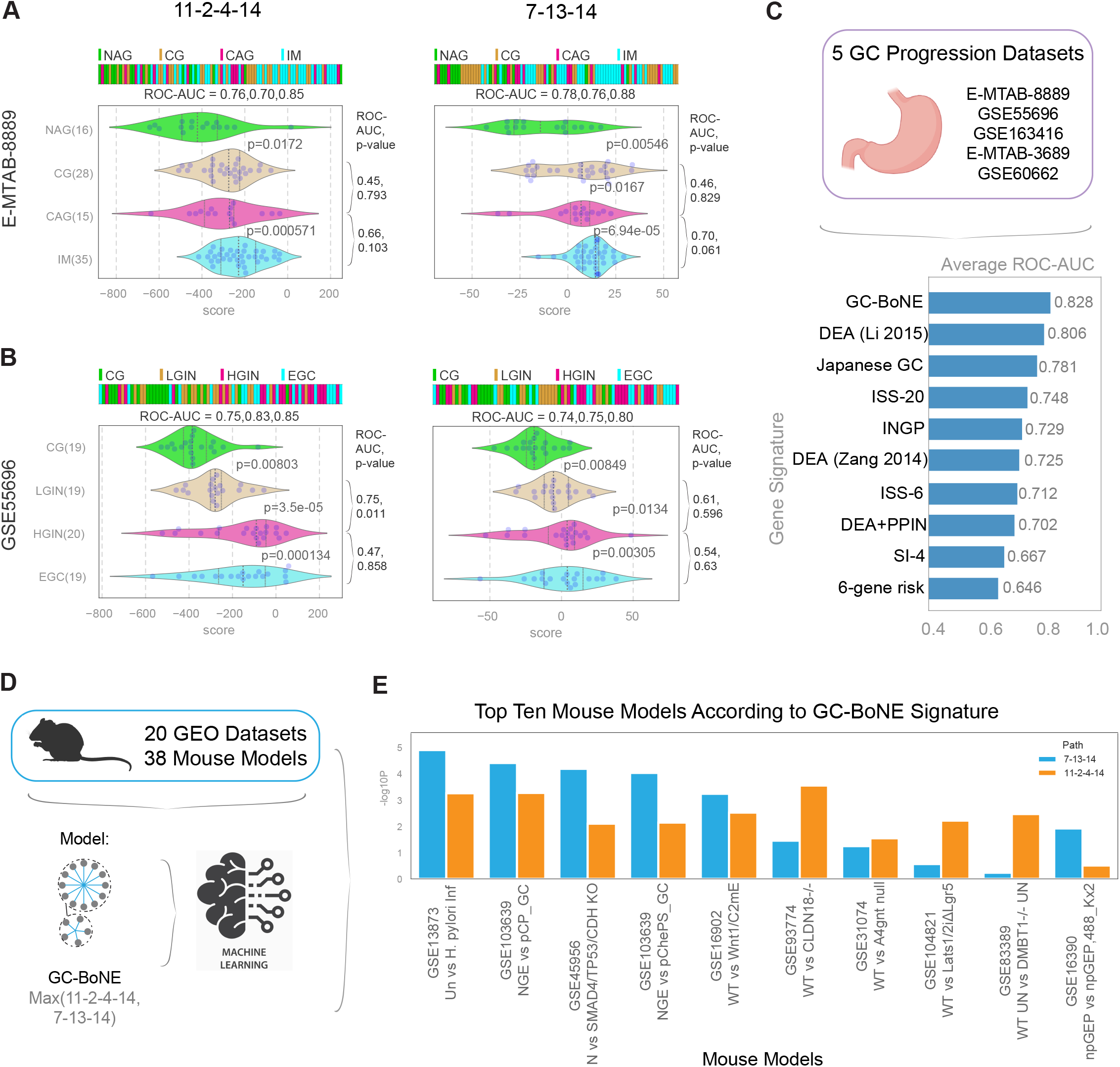
GC-BoNE signature in GC progression and mouse models. **a**. Violin plots for GC progression of normal active gastritis (NAG) → chronic active gastritis (CG) → chronic atrophic gastritis (CAG) → intestinal metaplasia (IM) (E-MTAB-8889) using the GC-BoNE signature: 11-2-4-14 (*left*) and 7-13-14 (*right*) **b**. Violin plots for chronic gastritis (CG) → low-grade intestinal neoplasia (LGIN) → high-grade intestinal neoplasia (HGIN) → early gastric cancer (EGC) (GSE55696) using the GC-BoNE signature: 11-2-4-14 (*left*) and 7-13-14 (*right*) **c**. Comparison of average ROC-AUC values for GC progression datasets using GC-BoNE and other gene signatures **d**. Schematic summarizing comparison of 38 mouse models from 20 GEO datasets using GC-BoNE **e**. Top ten mouse models according to –log_10_(p-value) from Welch’s Two Sample t-test separated by path (7-13-14: blue, 11-2-4-14: orange)

### GC-BoNE can objectively assess the appropriateness of mouse models for studying human GC

Next, we wanted to identify mouse models that recapitulated human normal versus GC. We analyzed 38 mouse models [24-41] from 20 NCBI GEO datasets using C#11-2-4-14 and C#7-13-14 (see **Online Resource 2** for a list of GSE IDs; **Fig. 3d**). Many of the mouse models had a perfect ROC-AUC of 1.00 using C#11-2-4-14 and C#7-13-14 (see **Online Resource 4**). We then looked at which mouse models are significantly different using a t-test to determine the top ten models (**Fig. 3e**). It is noteworthy that the top two models represent the two common risk factors for GC in humans. The model that ranked #1 (GSE13873) is one in which the *H. pylori* infection →GC cascade is modeled in C57Bl6 mouse model of experimental infection with the closely related *H. felis*. The authors showed that while most infected mice develop premalignant lesions such as gastric atrophy, compensatory epithelial hyperplasia and IM, a minority is completely protected from preneoplasia. The models that ranked #2-6 (GSE103639 (NGE vs pCP_GC), GSE45956, GSE103639 (NGE vs pChePS_GC), GSE16902, GSE93774) were all genetically engineered mouse models (GEMMs) in which targeted deletions were performed on genes (*CDH1, SMAD4, CLDN18* etc.) that are associated with risk of GC, by virtue of being either the most common germline mutation in GC (*CDH1 [42]*), or for harboring disease-associated SNPs (*SMAD4 [43]*) or being the target of the most frequent somatic genomic rearrangements [44] (*CLDN18*). These results suggest that *GC-BoNE* can objectively assess the degree of similarity between mouse models (both infection-induced and genetically-induced types) and human GC. In doing so, it can pinpoint which mouse models best recapitulate the patterns of gene expression that is observed during the transformation from healthy to GC in human samples.

### GC-BoNE (C#11-2-4-14) can prognosticate the risk of IM →GC progression

Because we want to identify genes responsible for the progression of GC, we looked at a dataset that curated samples from a prospective study [45] with long-term follow-up (a mean of 12±3.4 years) to evaluate risk of progression to GC among patients with incomplete or complete intestinal metaplasia (IIM and CIM respectively) (**Fig. 4a**). It is known that among the types of intestinal metaplasia, IIM carries a greater risk for progression to GC compared to CIM [46]. A recent meta-analysis showed that compared with CIM, pooled RR of cancer/dysplasia in IIM patients was 4.48 (95% CI 2.50–8.03), and the RR was 4.96 (95% CI 2.72–9.04) for cancer, and 4.82 (95% CI 1.45– 16.0) for dysplasia [47]. We found that C#11-2-4-14 best distinguished the healthy control patients (HC), patients with high risk-carrying IIM that progressed (IIM-GC) and those that did not progress (IIM-C) (ROC-AUC values: HC vs IIM-C: 0.86, HC vs IIM-GC: 0.94, IIM-C vs IIM-GC: 0.95; **Fig. 4b**). C#11-2-4-14 was not able to significantly distinguish (using Student’s t-test) low risk-carrying CIM from HC. C#7-13-14 also could distinguish HC vs IIM-C (ROC-AUC = 0.80) and HC vs IIM-GC (ROC-AUC = 0.88), but not IIM-C vs IIM-GC (ROC-AUC = 0.71), however C#11-2-4-14 performed better (**Fig. 4c**). The DEA (Li 2015) gene signature similarly separates HC vs IIM-C (ROC-AUC = 0.90) and HC vs IIM-GC (ROC-AUC = 0.87) but is not able to distinguish IIM-C vs IIM-GC (ROC-AUC = 0.38) (**Fig. 4d**). The Japanese GC signature cannot significantly distinguish any of the samples (ROC-AUC values range from 0.42 - 0.74; **Fig. 4e**). These findings suggest genes in C#11-2-4-14 might be key to understanding why some IIM patients progress to GC.

**Fig. 4.**
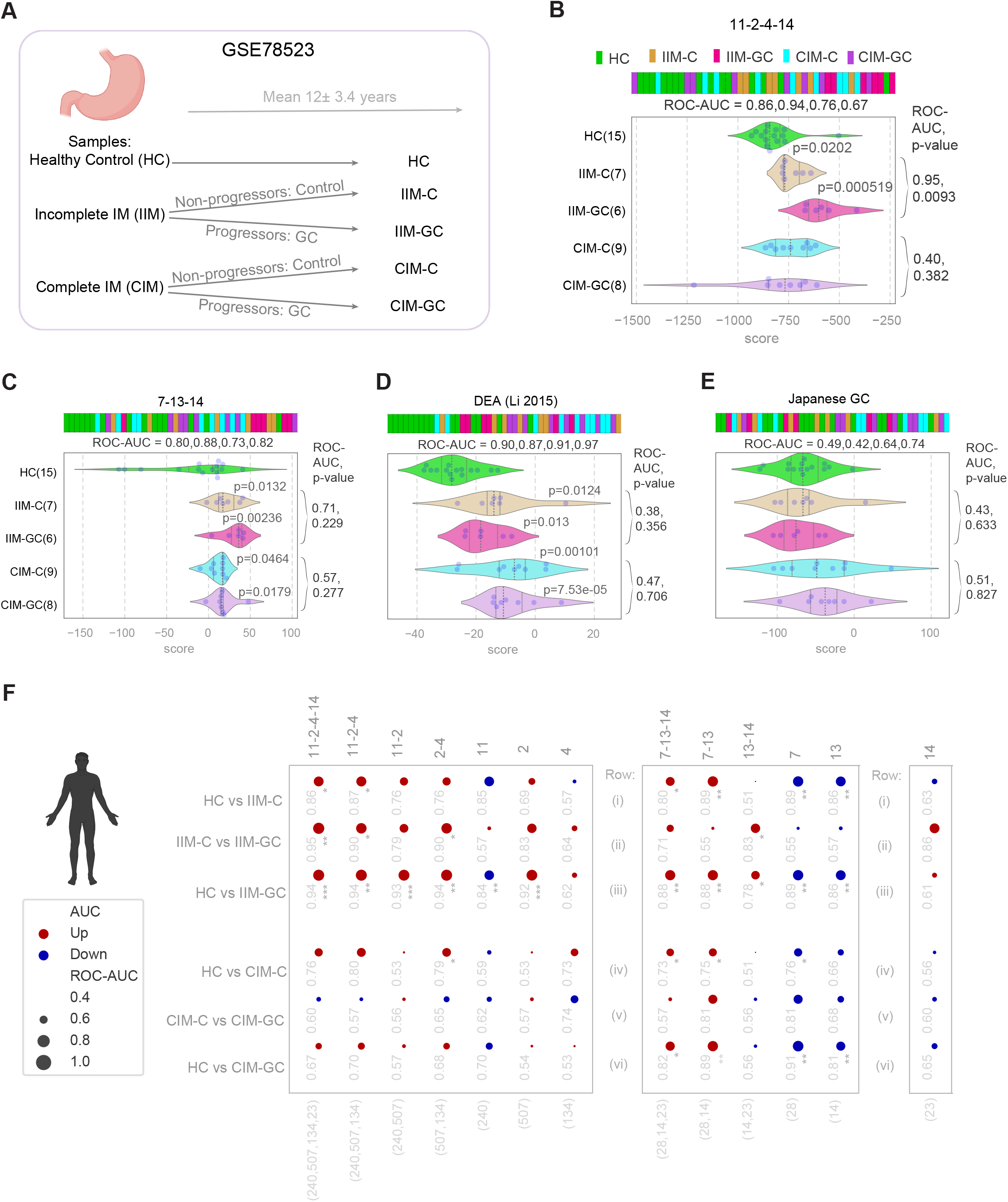
GC-BoNE signature predicts outcome. **a**. Schematic summarizing GSE78523: samples collected from healthy patients (HC) and patients with incomplete IM (IIM) or complete IM (CIM). After a mean of 12 += 3.4 years, patients with IM were diagnosed as non-progressors (control: C) or progressors (GC) **b-e**. Violin plots showing classification of samples using GC-BoNE, DEA (Li 2015), and Japanese GC signatures (**b**: 11-2-4-14, **c**: 7-13-14, **d**: DEA (Li 2015), **e**: Japanese GC) **f**. GSE78523 is visualized as bubble plots of ROC-AUC values (radius of circles is based on the ROC-AUC) demonstrating the direction of gene regulation (Up: red, Down: blue) for the classification of samples (GC-BoNE clusters in columns; sample comparison in rows). P-values based on Welch’s T-test (of composite score of gene expression values) are provided using the standard code (*p<=0.05, **p<=0.01, ***p<=0.001) next to the ROC-AUC

### GC-BoNE provides insights into the changes in cellular continuum states during healthy →IIM →GC progression

To understand which cellular processes change during cell transformation and which genes contribute to the progression of GC, we checked how clusters in C#11-2-4-14 and C#7-13-14 perform separately (**Fig. 4f**). When looking at HC vs IIM-C (**Fig. 4f row i**), cluster 14 is not able to distinguish the samples (ROC-AUC = 0.63), but both C#11-2-4 and C#7-13 are able to separate the samples (ROC-AUC = 0.87, 0.89 respectively). However, when you compare IIM-C vs IIM-GC (**Fig. 4f row ii**), cluster 14 is better able to distinguish the samples (ROC-AUC = 0.86), with C#11-2-4-14 best able to classify the samples (ROC-AUC = 0.95). These results show genes in C#11-2-4 might be responsible for the progression from HC to IIM, while C#14 is important for IIM to GC. Findings thereby suggest that the progression from HC to IIM may be impacted by genes related to muscle contraction, cell cycle and immune system, while the progression from IIM to GC is affected by extracellular matrix processes.

## DISCUSSION

Although the incidence rates of GC have been decreasing around the world [4], there have not been any significant improvements in terms of new therapeutics, diagnostics and changes in screening designed for preneoplastic stages. In this study, we built a Boolean implication network using GSE66229 and used machine learning (on GSE37023 and [GSE122401](https://www.ncbi.nlm.nih.gov/geo/query/acc.cgi?acc=GSE37023)) to identify a gene signature (GC-BoNE) which could classify normal and gastric samples. Reactome pathway analysis of GC-BoNE revealed the following biological processes in the GC tumor samples: decrease in muscle contraction and ion transport, and increase in cell cycle, immune system and extracellular matrix functions (**Fig. 1e**). Although previous studies have identified most of these pathways [15-22], muscle contraction has not been widely identified, providing a new area to focus on researching. We then tested how GC-BoNE compares to gene signatures from past studies in both normal vs GC samples (**Fig. 2c**) and GC progression samples (**Fig. 3c and 4f**).

Our Boolean network-based approach improves upon past studies by *<u>First</u>*, identifying a gene signature (GC-BoNE) that is better able to classify samples along the GC disease continuum compared to previous signatures. When looking at normal vs GC samples, many of the signatures performed well (**Fig. 2c**). However, we are more interested in finding a gene signature that can distinguish samples earlier in the GC disease continuum. When looking at GC progression, our signature outperforms the other gene signatures (**Fig. 3c**). Since the genes in GC-BoNE do not overlap with many genes from the other gene signatures (**Fig. 1e**), this provides a list of new potential biomarkers for targeting therapeutics at different points along the GC disease continuum.

*<u>Second</u>*, we identified C#11-2-4 as important in the progression for HC to IIM-C, while C#14 is important for the progression from IIM-C to IIM-GC (**Fig. 4f**). Although the model was built and trained on N vs GC samples, using a Boolean network-based approach allows us to identify paths that can also determine the intermediate states of disease progression. The invariant asymmetric Boolean implications present in the GC-BoNE signature provide insight into the cellular changes occurring at various time points along the disease continuum. These findings provide a list of gene targets that can be tested using the mouse models we identified (**Fig. 3e**) or other models. Genes in C#11-2-4 with cellular processes affecting muscle contraction, cell cycle and immune system can be targeted for drug development in patients with IIM before they advance to GC. Genes affecting extracellular matrix processes in C#14 can be targeted for patients with GC.

Overall, we demonstrate that the genes identified from our Boolean network-based approach were better able to classify samples along the GC disease continuum compared to the genes from previous work. The genes from GC-BoNE provide more opportunities to research the cellular processes behind GC progression. Results from this paper can be used to rationalize gene targets for diagnostics and therapeutics.

## AUTHORS’ DISCLOSURES

Authors declare that they have no competing interests.

## AUTHORS’ CONTRIBUTIONS

Conceptualization: D.S, P.G

Methodology: D.S, D.V.

Investigation: D.V, D.S, P.G

Visualization: D.V, D.S, P.G

Funding acquisition: D.S, P.G

Project administration: D.S, P.G

Supervision: D.S, P.G

Writing – original draft: D.V, D.S, P.G

Writing – review & editing: D.V, D.S, P.G

## ACKNOWLEDGEMENTS

This work was supported by the National Institutes for Health (NIH) grant R01-AI155696 (to PG and DS). Other sources of support include: T32GM139790 (to DV), R01-GM138385 (to DS), R01-AI141630, CA100768 and CA160911 (to P.G), and UG3TR002968 (to D.S. and P.G). D.S was also supported by two Padres Pedal the Cause awards (Padres Pedal the Cause/RADY #PTC2017 and San Diego NCI Cancer Centers Council (C3) #PTC2017). D.S and P.G were also supported by the Leona M. and Harry B. Helmsley Charitable Trust.

We would also like to thank Saptarshi Sinha, Dharanidhar Dang and Sahar Taheri for providing feedback on the manuscript.

## ONLINE RESOURCES

**Online Resource 1** Supplementary methods

**Online Resource 2** List of GSE IDs used in the analysis along with sample type (human vs mouse), use (network, training, validation) and figure panel

**Online Resource 3** Complete list of genes used in all gene signatures (GC-BoNE and signatures from other sources)

**Online Resource 4** Bubble plots of ROC-AUC values (radius of circles is based on the ROC-AUC) demonstrating the direction of gene regulation (Up: red, Down: blue) for the classification of samples in 38 mouse models (GC-BoNE clusters in columns; sample comparison in rows). P-values based on Welch’s T-test (of composite score of gene expression values) are provided using the standard code (*p<=0.05, **p<=0.01, ***p<=0.001) next to the ROC-AUC

## Notes

### Competing Interest Statement

The authors have declared no competing interest.

