## Supplementary Methods 1 for "Artificial Intelligence Guided Discovery of Gastric Cancer Continuum"

### Materials and Methods

#### Computational Approaches

*An AI-assisted study design that uses Boolean approach to build transcriptomic networks*

We chose a Boolean approach to building transcriptomic networks[1] because of its ability to pinpoint with precision cellular states in tissues. For example, it helped pinpoint branchpoints in B/T cell differentiation[2, 3], define progenitor cell hierarchy in blood[4-8], normal and neoplastic cell states in colorectal cancers[9, 10], bladder cancers[11-14], and prostate cancers[15, 16], and identify NK cell exhaustive states[17], universal cell proliferative[18] and macrophage[19] markers, and cell states in the mucosal barrier in IBD[20]. Because the Boolean approach relies on invariant relationships that are conserved despite heterogeneity in the samples used for the analysis, which often represent maximum possible diversity, i.e., the relationships can be thought of as general relationships among pairs of genes across all samples irrespective of their origin (normal or disease), laboratories/cohorts, different perturbations, and sometimes in multiple species including human, mouse and rat, and hence, considered conserved invariants. It is assumed that such 'invariants' are likely to be fundamentally important for any given process.

*Gastric cancer datasets used for network analysis*

One microarray dataset (GSE66229; n = 400, 300 gastric cancer (GC) tumor and 100 patient-matched normal tissue) is used to perform Boolean Implication network analysis of GC samples. We used the $log_2$ RMA normalized data instead of the $log_{10}$ RMA normalized data available on GEO. Two microarray datasets (GSE37023 (only samples on GPL96 Affymetrix Human Genome U133A Array used for analysis), n = 65, non-malignant = 36, GC tumor = 29; GSE122401, n = 160, patient-matched normal = 80, GC tumor = 80) are used to train a network model to distinguish normal vs GC samples. All training and validation dataset (**Supplementary Information 1**) were downloaded from National Center for Biotechnology Information (NCBI) Gene Expression Omnibus website (GEO)[21-23] or European Molecular Biology Laboratory (EMBL) European Bioinformatics Institute (EMBL-EBI) ArrayExpress website[24]. All gene expression datasets (**Supplementary Information 1**) were processed separately using the Hegemon data analysis framework[9, 10, 12].

*Boolean Analysis*

*Boolean logic* is a simple mathematical relationship of two values, i.e., high/low, 1/0, or positive/negative. The Boolean analysis of gene expression data requires first the conversion of expression levels into two possible values. The *StepMiner* algorithm is reused to perform Boolean analysis of gene expression data[1]. The *Boolean analysis* is a statistical approach which creates binary logical inferences that explain the relationships between phenomena. Boolean analysis is performed to determine the relationship between the expression levels of pairs of genes. The *StepMiner* algorithm is applied to gene expression levels to convert them into Boolean values (high and low). In this algorithm, first the expression values are sorted from low to high and a rising step function is fitted to the series to identify the threshold. Middle of the step is used as the *StepMiner* threshold. This threshold is used to convert gene expression values into Boolean values. A noise margin of 2-fold change is applied around the threshold to determine intermediate values, and these values are ignored during Boolean analysis. In a scatter plot, there are four possible quadrants based on Boolean values: (low, low), (low, high), (high, low), (high, high).

*Invariant Boolean implication relationships*

A Boolean implication relationship is observed if any one of the four possible quadrants or two diagonally opposite quadrants are sparsely populated. Based on this rule, there are six different kinds of Boolean implication relationships. Two of them are symmetric: equivalent (corresponding to the highly positively correlated genes), opposite (corresponding to the highly negatively correlated genes). Four of the Boolean relationships are asymmetric, and each corresponds to one sparse quadrant: (low => low), (high => low), (low => high), (high => high). BooleanNet statistics (Equations listed below) are used to assess the sparsity of a quadrant and the significance of the Boolean implication relationships[1, 2]. Given a pair of genes A and B, four quadrants are identified by using the StepMiner thresholds on A and B by ignoring the Intermediate values defined by the noise margin of 2-fold change (+/- 0.5 around StepMiner threshold). The number of samples in each quadrant are defined as $a_{00}$, $a_{01}$, $a_{10}$, and $a_{11}$. Total number of samples where gene expression values for A and B are low is computed using the following equations.

$$nA_{low} = (a_{00} + a_{01}),\ nB_{low} = (a_{00} + a_{10}),$$

The total number of samples considered is computed using the following equation.

$$total = a_{00} + a_{01} + a_{10} + a_{11}$$

Expected number of samples in each quadrant is computed by assuming independence between A and B. For example, expected number of samples in the bottom left quadrant $e_{00} = \hat{n}$ is computed as probability of A low (($a_{00}$ + $a_{01}$)/total) multiplied by probability of B low (($a_{00}$ + $a_{10}$)/total) multiplied by total number of samples. The following equation is used to compute the expected number of samples.

$$n = a_{ij}, \hat{n} = \left(\frac{nA_{low}}{total} * \frac{nB_{low}}{total}\right) * total$$

To check whether a quadrant is sparse, a statistical test for ($e_{00} > a_{00}$) or $(\hat{n} > n)$ is performed by computing $S_{00}$ and $p_{00}$ using the following equations. A quadrant is considered sparse if $S_{00}$ is high $(\hat{n} > n)$ and $p_{00}$ is small.

$$S_{ij} = \frac{\hat{n} - n}{\sqrt{\hat{n}}}$$

$$p_{00} = \frac{1}{2}(\frac{a_{00}}{(a_{00} + a_{01})} + \frac{a_{00}}{(a_{00} + a_{10})})$$

A threshold of $S_{00}$ > sthr and $p_{00}$ < pthr to check sparse quadrant. A Boolean implication relationship is identified when a sparse quadrant is discovered using the following equation.

***Boolean Implication*** = ($S_{ij}$ > sthr, $p_{ij}$ < pthr)

A relationship is called Boolean equivalent if top-left and bottom-right quadrants are sparse.

*Equivalent* $= (S_{01} > sthr, P_{01} < pthr, S_{10} > sthr, P_{10} < pthr$

Boolean opposite relationships have sparse top-right ($a_{11}$) and bottom-left ($a_{00}$) quadrants.

$Opposite = (S_{00} > sthr, P_{00} < pthr, S_{11} > sthr, P_{11} < pthr$

Boolean equivalent and opposite are symmetric relationship because the relationship from A to B is same as from B to A. Asymmetric relationship forms when there is only one quadrant sparse (A low => B low: top-left; A low => B high: bottom-left; A high=> B high: bottom-right; A high => B low: top-right). These relationships are asymmetric because the relationship from A to B is different from B to A. For example, A low => B low and B low => A low are two different relationships.

A low => B high is discovered if bottom-left ($a_{00}$) quadrant is sparse and this relationship satisfies following conditions.

*A low => B high* = $(S_{00} > sthr, P_{00} < pthr)$

Similarly, A low => B low is identified if top-left ($a_{01}$) quadrant is sparse.

*A low => B low* = $(S_{01} > sthr, P_{01} < pthr)$

A high => B high Boolean implication is established if bottom-right ($a_{10}$) quadrant is sparse as described below.

*A high => B high* = $(S_{10} > sthr, P_{10} < pthr)$

Boolean implication A high => B low is found if top-right ($a_{11}$) quadrant is sparse using following equation.

*A high => B low* = $(S_{11} > sthr, P_{11} < pthr)$

For each quadrant, a statistic $S_{ij}$ and an error rate $p_{ij}$ is computed. $S_{ij} > 3$ and $p_{ij} < 0.1$ are the thresholds used on the BooleanNet statistics to identify Boolean implication relationships (BIRs). False discovery rate is computed by randomly shuffling each gene and computing the ratio of the number of Boolean implication relationship discovered in the randomized dataset and original dataset. The false discovery rate for GC dataset was less than 0.001.

Boolean Implication analysis looks for invariant relationships across all the different types of samples regardless of the conditions and treatment protocols. Therefore, it does not distinguish the sample types when discovering Boolean implication relationships. We assume there are fundamental invariant Boolean implication formulas that are satisfied by every sample regardless of their type.

*Construction of GC Boolean Implication Network*

A Boolean implication network (BIN) is created by identifying all significant pairwise Boolean implication relationships (BIRs)[24, 25]. The Boolean implication network contains the six possible Boolean relationships between genes in the form of a directed graph with nodes as genes and edges as the Boolean relationship between the genes. The nodes in the BIN are genes and the edges correspond to BIRs. Equivalent and Opposite relationships are denoted by undirected edges and the other four types (low => low; high => low; low => high; high => high) of BIRs are denoted by having a directed edge between them. The network of equivalences seems to follow a scale-free trend; however, other asymmetric relations in the network do not follow scale-free properties. BIR is strong and robust when the sample sizes are usually more than 200 (from our experience of using Boolean Implication for more than 10 years). All our previous papers used thousands of diverse samples to establish Boolean implication relationships. Boolean Implication analysis is carried out in GSE66229 (n = 400, 300 gastric cancer (GC) tumor and 100 patient-matched normal tissue) which has reasonable number of samples. We have demonstrated that we have excellent False Discovery Rate (< 0.001) when S > 3 and p < 0.1 are used. GSE66229 was prepared for Boolean analysis by filtering genes that had a reasonable dynamic range of expression values. For each gene, probesets with best dynamic range was chosen. When the dynamic range of expression values was small, it was difficult to distinguish if the values were all low or all high or there were some high and some low values. Thus, it was determined to be best to ignore them during Boolean analysis. The filtering step was performed by analyzing the fraction of high and low values identified by the StepMiner algorithm[26]. Any probe set or genes which contained less than 5% of high or low values were dropped from the analysis.

*Generation of Clustered Boolean Implication network*

Clustering was performed in the Boolean implication network to dramatically reduce the complexity of the network. A clustered Boolean implication network (CBIN) was created by clustering nodes in the original BIN by following the equivalent BIRs. One approach is to build connected components in an undirected graph of Boolean equivalences. However, because of noise, the connected components become internally inconsistent e.g., two genes opposite to each other become part of the same connected component. In addition, the size of clusters became unusually big with almost everything in one cluster. To avoid such a situation, we need to break the component by removing the weak links. To identify the weakest links, we first computed a minimum spanning tree for the graph and computed the Jaccard similarity coefficient for every edge in this tree. Ideally if two members are part of the same cluster, they should share as many connections as possible. A threshold is considered for the Jaccard similarity coefficient (0.5 for GC network) below which the edges are dropped from further analysis. Thus, many weak equivalences were dropped using the above algorithm leaving the clusters internally consistent. We removed all edges that have Jaccard similarity coefficient less than the selected threshold and built the connected components with the rest. The connected components were used to cluster the BIN which is converted to the nodes of the CBIN. The choice of the threshold on the Jaccard similarity coefficient plays a significant role in determining the size and the number of clusters as well as whether they are internally consistent. A new graph was built that connected the individual clusters to each other using Boolean relationships. The link between two clusters (A, B) was established by using the top representative node from A that was connected to most of the members of A and sampling 6 nodes from cluster B and identifying the overwhelming majority of BIRs between the nodes from each cluster.

A CBIN was created using the GSE66229 dataset. Each cluster was associated with healthy or disease samples based on where these gene clusters are highly expressed. The edges between the clusters represented the Boolean relationships that are color-coded as follows: orange for low => high, dark blue for low => low, green for high => high, red for high => low, light blue for the equivalent and black for the opposite.

*Charting Boolean paths*

Boolean paths have been explored before to predict the underlying time series events in biological processes such as B cell differentiation[2, 3] and early differentiation events in cancer stem cells[9, 10, 12, 15]. This algorithm is called MiDReG (Mining Developmentally Regulated Genes) that uses two seed genes to identify intermediate genes in a biological process. MiDReG infers intermediate states using a sequence of asymmetric BIRs. Here, using MiDReg algorithm/concept to traverse the Boolean Implication network that identifies paths of clusters where the start and end clusters in the clustered Boolean implication network mark the end points of a possible set of events from healthy to disease. The asymmetric BIRs provide a unique dimension to the network that is fundamentally different from any other gene expression networks in the literature. Traversing a set of nodes in a directed graph of the Boolean network constitutes a Boolean path. A simple Boolean path involves two nodes and the directed edge between them. A complex Boolean path involves more than two nodes and the edges between them.

*Ordering samples based on composite score of Boolean path*

A Boolean path contains one or more clusters. A composite score is computed for each cluster and combined later. To compute the final score, first the genes present in each cluster were normalized and averaged. Gene expression values were normalized according to a modified Z-score approach centered around StepMiner threshold (formula = (expr - SThr)/3/stddev). A weighted linear combination of the averages from the clusters of a Boolean path was used to create a score for each sample. The weights along the path either monotonically increased or decreased to make the sample order consistent with the logical order based on BIR. The samples were ordered based on the final weighted and linearly combined score. A cluster highly expressed in a disease setting received a positive weight (ex: 1, 2, 3, etc.) and healthy setting received a negative weight (ex: -1, -2, -3, etc.).

*Summary of genes in the clusters*

Reactome pathway analysis of each cluster along the top continuum paths was performed to identify the enriched pathways[27]. The pathway description was used to summarize at a high-level what kind of biological processes are enriched in a particular cluster.

*Measurement of classification strength or prediction accuracy*

Receiver operating characteristic (ROC) curves were computed by simulating a score based on the ordering of samples that illustrates the diagnostic ability of binary classifier system as its discrimination threshold is varied along with the sample order. The ROC curves were created by plotting the true positive rate (TPR) against the false positive rate (FPR) at various threshold settings. The area under the curve (often referred to as simply the AUC) is equal to the probability that a classifier will rank a randomly chosen IBD samples higher than a randomly chosen healthy samples. In addition to ROC AUC, other classification metrics such as accuracy ((TP + TN)/N; TP: True Positive; TN: True Negative; N: Total Number), precision (TP/(TP+FP); FP: False Positive), recall (TP/(TP+FN); FN: False Negative) and f1 (2 * (precision * recall)/(precision + recall)) scores were computed. Precision score represents how many selected items are relevant and recall score represents how many relevant items are selected. Fisher exact test is used to examine the significance of the association (contingency) between two different classification systems (one of them can be ground truth as a reference).

*AI guided discovery of Boolean paths*

A Boolean path is converted to a path score as mentioned above using a linear combination of normalized gene expression values. The strength of classification of normal and GC samples using this score is computed by the ROC-AUC measurement. We performed multivariate regression to identify the best Boolean path that predicts AN vs GC in GSE37023 and GSE122401. We tested how the path score distinguished normal/AN, GC tumor and GC progression samples as they are annotated in many other independent datasets.

*Training and Validation Datasets*

A Boolean path is selected after machine learning to construct a Boolean model. The Boolean model is tested in several human and mouse datasets, each comprised of a heterogeneous collection of samples (as mentioned in **Supplementary Information 1**) to demonstrate reproducibility. Selected Boolean path score is computed as mentioned in section "*Ordering samples based on composite score of Boolean path*". The sample order using the Boolean path score is evaluated using the sample annotation (N vs T, AN vs T, and N+AN vs T) by ROC-AUC analysis. We tested how the 11-2-4-14 path score with weight -1, 1, 2, 4 respectively and 7-13-14 with weight -2, -1, 2 distinguishes normal and GC samples as they are annotated in training datasets (GSE37023 and GSE122401), and validation datasets (GSE54129, GSE2669, GSE33651, GSE2637, GSE65801, TCGA-STAD, GSE27342, GSE118916, E-MTAB-6693, GSE29272, GSE13861, GSE38940, GSE31811, GSE122796, GSE13911, GSE29998, GSE33335, GSE2138, E-MTAB-9990, GSE9973, GSE19826). We also tested these paths in GC progression datasets (E-MTAB-8889, GSE55696, GSE163416, E-MTAB-3689, GSE60662, GSE78523). We have collected publicly available gene expression datasets derived from mouse models of GC (GSE13873, GSE69145, GSE69146, GSE166904, GSE83389, GSE142644, GSE104821, GSE16390, GSE102297, GSE43145, GSE27711, GSE145583, GSE93173, GSE31074, GSE164807, GSE144386, GSE93774, GSE103639, GSE45956, GSE16902; **Supplementary Data 1**) to test whether human Boolean models perform well in mice. The gene name conversion from human to mouse is performed using human genome GRCh38.95 ensembl IDs and mapping data exported from ensemble BioMart web-interface.

*Statistical Analysis*

Statistical significance between experimental groups was determined using Welch's Two Sample t-test (two-tailed, unpaired, unequal variance (equal_var=False), and unequal sample size). For all tests, a p-value of 0.05 was used as the cutoff to determine significance. All statistical analysis was performed using SciPy 1.5.4.
