## Supplementary figures and images for "Artificial Intelligence Guided Discovery of Gastric Cancer Continuum"

### Supplemental Figure 4

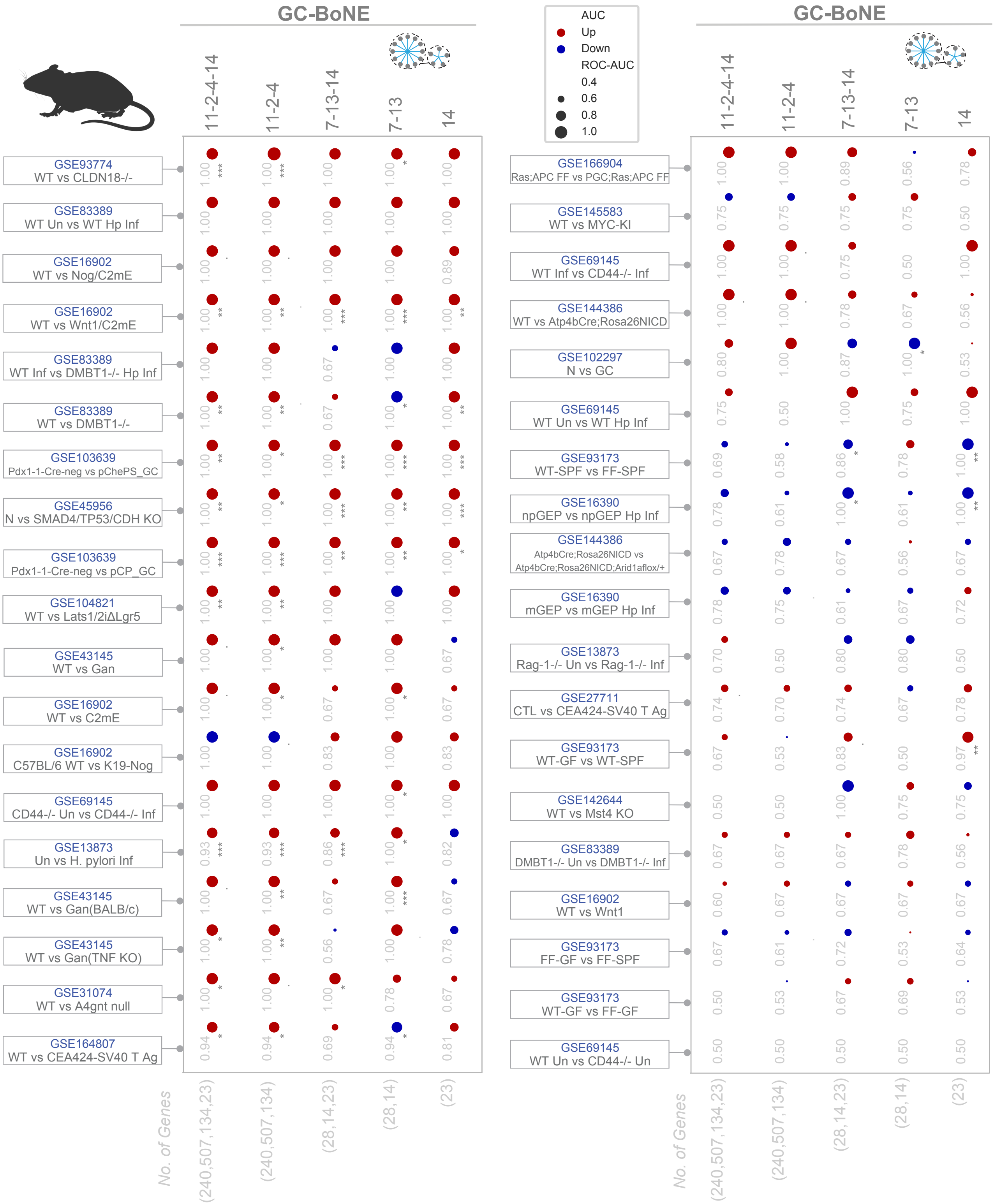
